# Human light meromyosin mutations linked to skeletal myopathies disrupt the coiled coil structure and myosin head sequestration

**DOI:** 10.1101/2023.05.15.540775

**Authors:** Glenn Carrington, Abbi Hau, Sarah Kosta, Hannah F. Dugdale, Francesco Muntoni, Adele D’Amico, Peter Van den Bergh, Norma B. Romero, Edoardo Malfatti, Juan Jesus Vilchez, Anders Oldfors, Sander Pajusalu, Katrin Õunap, Marta Giralt-Pujol, Edmar Zanoteli, Kenneth S. Campbell, Hiroyuki Iwamoto, Michelle Peckham, Julien Ochala

**Affiliations:** The Astbury Centre for Structural and Molecular Biology, Faculty of Biological Sciences, University of Leeds, Leeds, UK; School of Molecular and Cellular Biology, Faculty of Biological Sciences, University of Leeds, Leeds, UK; Centre of Human and Applied Physiological Sciences, School of Basic and Medical Biosciences, Faculty of Life Sciences & Medicine, King’s College London, UK; Randall Centre for Cell and Molecular Biophysics, School of Basic & Medical Biosciences, Faculty of Life Sciences & Medicine, King’s College London, UK; Department of Physiology, University of Kentucky, Lexington, Kentucky, USA; School of Sport, Exercise and Health Sciences, Loughborough University, Loughborough, UK; UCL Great Ormond Street Institute of Child Health, London, UK; NIHR Biomedical Research Centre at Great Ormond Street Hospital, Great Ormond Street, London, UK; Department of Neurosciences, Unit of Neuromuscular and Neurodegenerative Disorders, IRCCS Bambino Gesù Children’s Hospital, Rome, Italy; Neuromuscular Reference Center, Neurology Department, University Hospital Saint-Luc, Brussels, Belgium; Neuromuscular Morphology Unit, Institute of Myology, Myology Research Centre INSERM, Sorbonne University, Hôpital Pitié-Salpêtrière, Paris, France; APHP, Centre de Référence de Pathologie Neuromusculaire Nord-Est-Ile-de-France, Henri Mondor Hospital, Inserm U955, France; U1179 UVSQ-INSERM Handicap Neuromusculaire: Physiologie, Biothérapie et Pharmacologie appliquées, UFR Simone Veil-Santé, Université Versailles Saint Quentin en Yvelines, Paris-Saclay, France; Neuromuscular and Ataxias Research Group, Instituto de Investigación Sanitaria La Fe, Valencia, Spain; Centro de Investigación Biomédica en Red de Enfermedades Raras (CIBERER) Spain, Valencia, Spain; Department of Laboratory Medicine, University of Gothenburg, Gothenburg, Sweden; Genetics and Personalized Medicine Clinic, Tartu University Hospital, Tartu, Estonia; Department of Clinical Genetics, Institute of Clinical Medicine, University of Tartu, Tartu, Estonia; Universidade de São Paulo, Hospital das Clínicas, Faculdade de Medicina, Departamento de Neurologia, São Paulo SP, Brazil; Universidade Federal de São Paulo, Escola Paulista de Medicina, Departamento de Neurologia, São Paulo SP, Brazil; Division of Cardiovascular Medicine, University of Kentucky, Lexington, Kentucky, USA; SPring-8, Japan Synchrotron Radiation Research Institute, Hyogo, Japan; Department of Biomedical Sciences, University of Copenhagen, Copenhagen, Denmark

**Keywords:** **<u>Keywords</u>**: Skeletal muscle, Laing distal myopathy, myosin storage myopathy, myosin, contractility

## Abstract

Myosin heavy chains encoded by *MYH7* and *MYH2* are among the most abundant proteins in human skeletal muscle. After decades of intense research using a wide range of biophysical and biological approaches, their functions have begun to be elucidated. Despite this, it remains unclear how mutations in these genes and resultant proteins disrupt myosin structure and function, inducing pathological states and skeletal myopathies termed myosinopathies. Here, we have analysed the effects of several common *MYH7* and *MYH2* mutations located in light meromyosin (LMM) using a broad range of approaches. We determined the secondary structure and filament forming capabilities of expressed and purified LMM constructs in vitro, performed *in-silico* modelling of LMM constructs, and evaluated the incorporation of eGFP-myosin heavy chain constructs into sarcomeres in cultured myotubes. Using muscle biopsies from patients, we applied Mant-ATP chase protocols to estimate the proportion of myosin heads that were super-relaxed, X-ray diffraction measurements to estimate myosin head order and myofibre mechanics to investigate contractile function. We found that human *MYH7* and *MYH2* LMM mutations commonly disrupt myosin coiled-coil structure and packing of filaments *in vitro*; decrease the myosin super-relaxed state *in vivo* and increase the basal myosin ATP consumption; but are not associated with myofibre contractile deficits. Altogether, these findings indicate that the structural remodelling resulting from LMM mutations induces a pathogenic state in which formation of shutdown heads is impaired, thus increasing myosin head ATP demand in the filaments, rather than affecting contractility. These key findings will help in the design of future therapies for myosinopathies.

## Introduction

Myosin, organised into thick filaments in skeletal and cardiac muscle, interacts with actin in the thin filaments to generate contraction and shortening. Muscle myosins are comprised of two heavy chains and two pairs of light chains (1). The first 838 residues (N-terminal region) form the myosin head, comprised of the motor and light chain binding domain (Fig. 1A). The motor domain interacts with actin and hydrolyses ATP. The light chain binding domain contains two IQ (isoleucine glutamine) repeats, of which the first binds essential light chain, and the second binds regulatory light chain (1). The downstream C-terminal sequence, starting after the invariant proline residue at position 838 (∼60%) forms an α-helix that dimerises with a second myosin heavy chain to form an α-helical coiled coil tail (Fig. 1B). The first third of the tail is known as subfragment-2 (S2), and the remaining C-terminal two thirds is known as light meromyosin (LMM).

**Figure 1.**
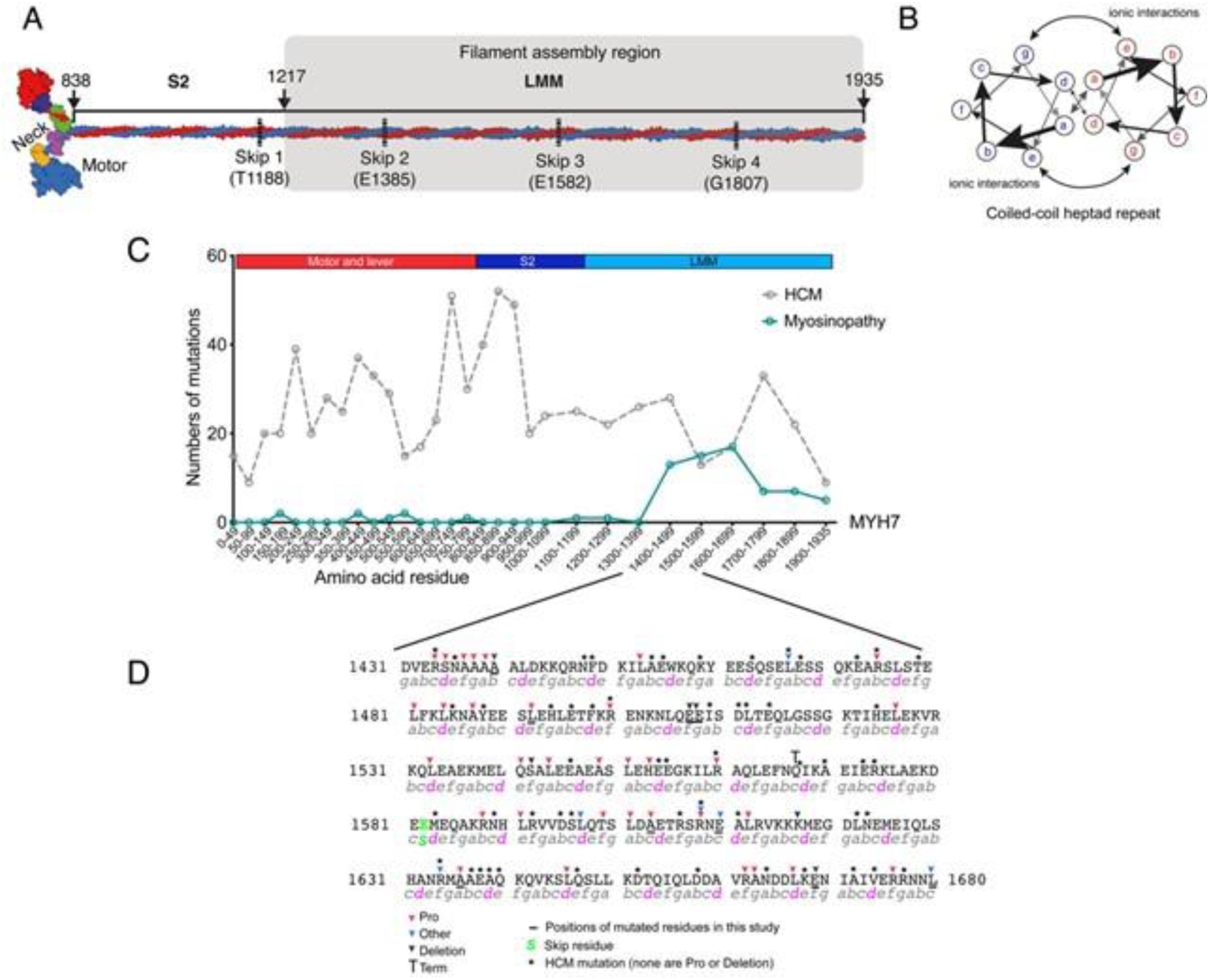
Myosinopathies and location of mutations. (A) A schematic showing the overall composition of striated myosin. The molecule is formed by two heavy chains that dimerise to form a coiled coil tail composed ofsubfragment-2 (S-2) and light-meromyosin (LMM)). The two heavy chains diverge to form the neck of lever, to which light chains bind, and the two motor domains, which bind actin and nucleotide. (B) is a schematic showing an end on view of the heptad repeat of two interacting α-helices. Residues in *a* and *d* positions form the hydrophobic seam. (C) displays the frequency of mutations in *MYH7* for Hypertrophic Cardiomyopathy (HCM) (grey dotted line) and for skeletal myopathies (Myosinopathy: green line), across the sequence of *MYH7*. (D) shows the sequence in which mutations that cause myosinopathies are most frequent. Positions of mutations (commonly mutation to proline or a single amino acid deletion) are indicated by coloured arrows. Mutations in residues mutated in HCM are indicated by *. Underlined residues indicate the position of mutated residues studied here. The heptad repeat is shown underneath the sequence, with *d* positions highlighted in magenta.

The LMM region of the myosin tail directs myosin assembly into filaments via clusters of alternating positive and negatively charge residues, each 28 residues long (42 Å) (2). This results in a quasi helical array of myosin heads on the surface of the filament, in which three pairs or heads protrude from the filament every 14.3 Å, forming a ‘crown’ of heads (3). This repeating feature gives rise to the M3 layer line in X-ray diffraction. The helical repeat, comprising three crowns, is 430 Å, and is equivalent to the axial stagger between adjacent LMM molecules in the thick filament. Two recent CryoEM structures of the C-zone (myosin-binding protein-C containing region) of cardiac muscle in relaxing conditions, have revealed the complex packing of the myosin tails (S2 and LMM) into the thick filament, together with the interactions of both myosin binding protein C (MyBPC) and titin with myosin in the filament, but not with each other (4, 5).

Until recently, striated muscle contraction was thought to be solely regulated by the binding of Ca^2+^ to thin filament proteins, shifting the position of tropomyosin on actin to reveal sites to which myosin prefers to bind (6). It has now become clear that activation of myosin heads, particularly in the C-zone of the thick filament is also important. In relaxed muscle, the myosin heads adopt a shutdown state in which the two heads interact with each other and with S2 to form an interacting heads motif (IHM) (7). This led to the idea that the heads need to be released from this state to drive contraction (8, 9). X-ray diffraction approaches have demonstrated that this release is indeed an important component of muscle contraction (10–12). Moreover, this shutdown state is likely to be linked to the super-relaxed state (SRX) state of myosin (8, 9) in which the ATPase rate is approximately 5 to 10 fold lower than that of disordered-relaxed (DRX) myosin heads (8, 13). The adoption of the SRX state is likely to be important in conserving energy between muscle contractions (8).

The key role of the SRX state in striated muscle has driven new concepts about how some mutations in myosin, and other striated muscle proteins might lead to striated muscle disease. A dominant missense mutation in the myosin heavy chain gene, *MYH7,* was first identified many years ago to cause hypertrophic cardiomyopathy (14). *MYH7* encodes the β/slow myosin heavy chain, which is expressed in both the heart and in slow skeletal muscle fibres. Over 1,000 missense mutations have since been described for *MYH7*, most of which are dominant, and these have now been implicated in several forms of heart disease (15). Approximately 50% of the known mutations are found in the myosin head, 20% in S2 and the remainder in LMM. While some mutations in the head are likely to directly affect the interaction with actin, others have been suggested to disrupt the IHM, shifting myosin heads into the DRX state, and thus increasing ATP consumption and metabolic demands on the muscle (9). Indeed, mavacamten, thought to promote the SRX state (16) has recently been approved as a drug to treat hypertrophic cardiomyopathy (17).

While the effect of mutations in the myosin head are becoming clearer, we still lack a clear understanding as to how the mutations in LMM result in muscle diseases. A subset of these mutations (about 70) in *MYH7* primarily result in skeletal muscle diseases known as myosinopathies rather than cardiomyopathies (18). The overall prevalence of myosinopathies is thought to be 1:26,000 (18) but this number is likely to be an under-estimate because of under-diagnosis (19, 20). Strikingly, almost all of these mutations are located in LMM (15). In addition, over 20 dominant/recessive mutations have recently been reported for *MYH2,* which encodes fast skeletal myosin heavy chain 2A, of which at least five are present in LMM (15).

The molecular and cellular mechanisms by which specific mutations in LMM lead to myosinopathies remain unclear and need addressing. We recently showed that two of these mutations, A1603P and K1617del, disrupt the structure of the coiled-coil and/or myosin filament formation (21) using a combination of in vitro and cellular assays. While LMM is not directly involved in stabilising the SRX state of myosin, disruption of myosin packing within the thick filament, with potential effects on titin and MyBPC organisation, could indirectly induce a decrease in heads in the SRX state, increasing ATP usage in skeletal muscle. Hence, in the present study, we performed a comprehensive analysis of several mutations in *MYH7* and *MYH2*, implicated in myosinopathies. We used circular dichroism and *in silico* modelling to determine the effects of mutations on secondary structure of LMM, EM to determine effects on LMM filament assembly, GFP-tagged myosin constructs to determine filament formation in cultured muscle cells, Mant-ATP assays to determine the ratio of SRX to DRX heads in samples from human tissue, as well as force and X-ray diffraction measurements of human tissue samples to gain a better understanding of how mutations in LMM cause myosinopathies.

## Results

### Mutations that cause myosinopathies are clustered in LMM

An analysis of mutations in *MYH7* that result in hypertrophic cardiomyopathy (HCM) and myosinopathies shows that mutations that cause HCM are distributed throughout the molecule, whereas those causing myosinopathies are mainly restricted to a specific region within LMM, between residues 1400-1700 (Fig. 1C). A closer look at the types of mutation that cause myosinopathies shows that they mainly comprise of mutations to proline (red arrows, Fig. 1D). The next most common type of mutation is a deletion of a single residue, while mutations to other residues are least common (Fig. 1D). In contrast, mutations that cause HCM, also found in this region, typically occur in different residues, are all missense mutations excluding mutation to proline, or single residue deletions (6). Proline residues are not usually found in coiled coils, as the kink that they introduce into the α-helix is typically not compatible with coiled coil formation. Deletion of a single residue introduces a shift in the heptad repeat pattern and is also likely to introduce a local disruption within the coiled coil. Thus, both types of mutation, found in myosinopathies, are likely to disrupt the secondary structure of the coiled coil, which in turn, could have effects on packing of LMM into filaments.

### LMM mutations alter the secondary structure of the coiled-coil

To determine the effects of specific mutations in LMM that result in myosinopathies, we measured the circular dichroism spectra of expressed and purified LMM constructs for MYH7 and MYH2. The constructs are expressed and purified as GST-LMM proteins, in which the two α-helices have already formed a coiled coil during expression in *E.coli*, and thus each of the α-helices contains the mutation. The GST is cleaved off prior to carrying out CD experiments.

These data showed that all the constructs were helical (Fig. 2A&D). However, the majority of the mutant constructs were less helical than the wild type constructs as measured by mean residue ellipticity (MRE) at 222 nm (Fig. 2C&D). Helicity was significantly decreased for all the mutations with the exception of MYH7 E1610K (Fig. 2C&D). Thus, either mutation to proline, or deletion does disrupt the secondary structure of the coiled coil.

**Figure 2.**
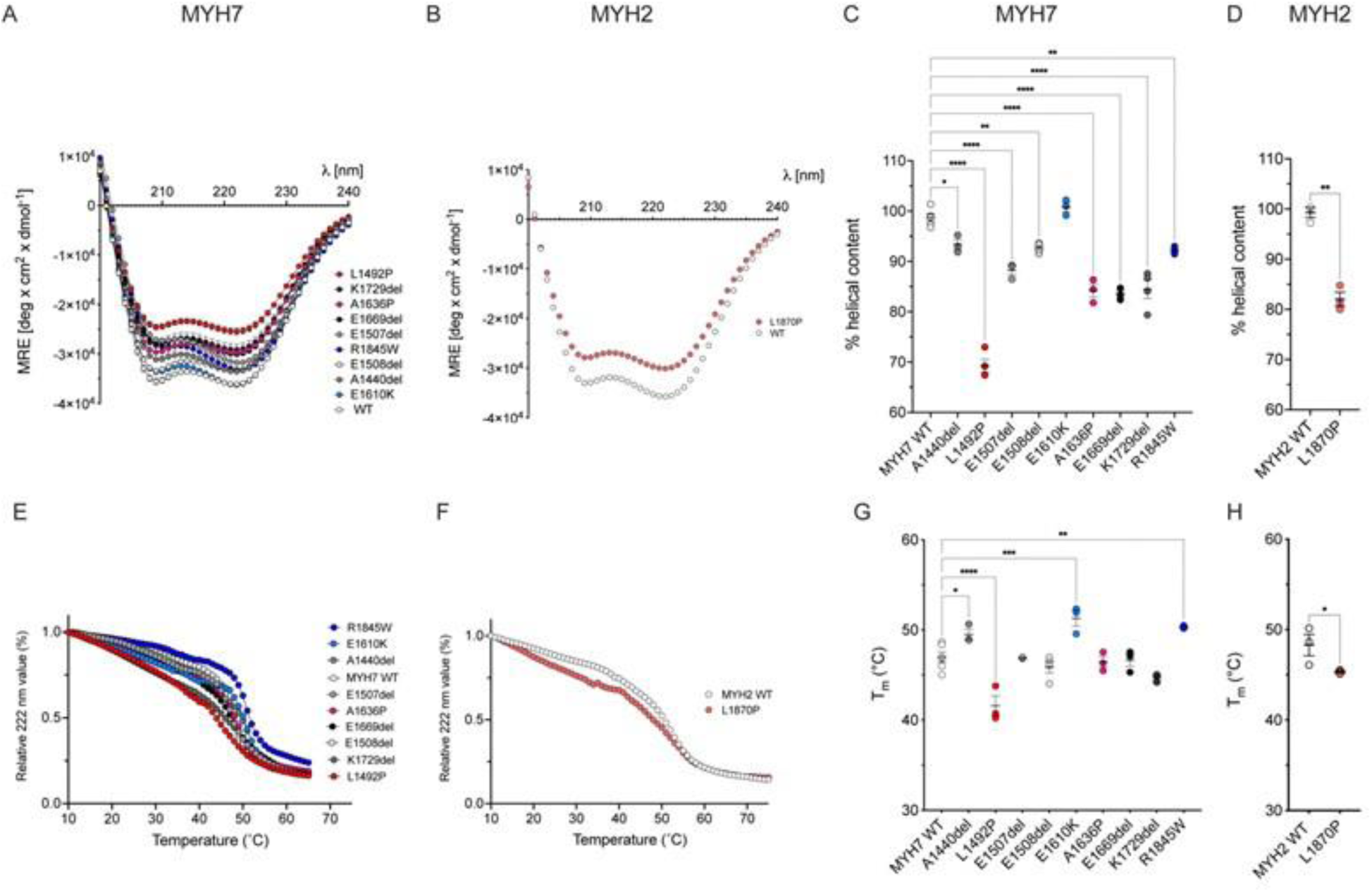
Secondary structure of LMM constructs. Circular dichroism spectra at 10°C for each of the *MYH7* (A) and *MYH2* (B) LMM constructs as shown in (A). Mutations to proline are shown in shades of red, deletion mutations in shades of grey and other mutations in shades of blue. The 222nm MRE values were used to estimate % helicity for each *MYH7* and *MYH2* construct (C & D). (E) and (F) show the thermal melt traces for MYH7 (E) and MYH2 (F) constructs. The 222 nm values were used to calculate Tm (temperature at which half the protein is melted) for each of the constructs (G & H). Individual data points, means and SEM are shown. Significant differences compared with WT are indicated; \**p* < 0.05; ** p<0.01; *** p<0.001; *** p<0.0001. Heptad positions for the mutated residues in MYH7: A1440 and A1636 ‘*b’;* E1610 and K1729 *‘c’;* L1492 *‘e’;* E1507 and R1845 *‘f’;* E1508 *‘g’* and in *MYH2: L1870 ‘d’*.

We then assessed the thermal stability of each of the mutations by evaluating the change in the MRE at 222 nm from 10 to 80°C and by calculating the temperature at which half the protein melted (Tm). Four of the nine MYH7 mutations tested here displayed changes in thermal stability. L1492P, which had the largest decrease in helicity (Fig. 2C) also had the largest decrease in thermal stability (Fig. 2G). K1729del, which had strongly reduced helical content, had a slightly lower thermal stability, but the decrease was not significant. In contrast, E1610K, which had a similar helical content to WT (Fig. 2C) had an increased thermal stability (Fig. 2G). The remaining MYH7 mutations showed no clear correlation between helical content and thermal stability. The helical content and thermal stability for the MYH2 mutation, L1870P were both significantly reduced.

To better characterise local perturbations to the structure of the coiled coil that could arise from each of the mutations, we performed molecular dynamics simulations similar to those we did previously (21). This allowed us to determine the local distances between the two helices (D_com_), the local heptad length and angle between adjacent heptad sections in each helix (Supplementary Fig. 1). In this approach, shorter wild type constructs were used either based on existing crystal structures or *de novo* models created using BEAMMOTIFCC (22). All the deletion mutants had a strong effect on Dcom, showing clear evidence for separation of both helices.

For the five deletion mutants in MYH7, the two helical strands tended to separate just after the mutation, except for E1669del, where there was strand separation before and after the mutation (Fig. S1A: Dcom). Each of these mutations showed effects of local heptad length close to the site of the mutation (Fig. S1B) and a pronounced effect on heptad angle (Fig. S1C). Thus, each of the deletion mutants had a considerable local disruption to the structure, introducing large local ‘kinks’ in the coiled-coil tail in order to accommodate the break in the heptad repeat and bring the coil back into register. This is consistent with the reduced helicity for each of these mutants in LMM, measured experimentally.

For the MYH7 proline mutation, L1492P, the strand separation around the site of the mutation was relatively small and there was a more pronounced increase in heptad length and a decrease in heptad angle (Fig. S1A-C). For A1636P, there was an increase in strand separation around the site of the mutation, but little effect on heptad length or angle. These differences may help explain the larger reduction in helicity and thermal stability experimentally observed for A1492P than for A1636P. For E1610K and R1845W, there was little strand separation around the site of the mutation (Fig. S1A), little change in heptad length (Fig S1B) and little change in heptad angle (Fig. S1C), consistent with small or no effects on helicity for these two mutants.

Finally, the MYH2 proline mutation, L1870P, showed a small increase in strand separation, a more pronounced increase in heptad length, and a decrease in heptad angle (Fig. S1D), somewhat similar to that observed for L1492P. This is consistent with the decreased helicity and thermal stability measured for this mutant.

### LMM mutations affect GST-LMM filament formation *in vitro* but incorporate normally into sarcomeres *in vivo*

GST-LMM molecules form short filaments *in vitro* in low ionic strength conditions (Fig. 3A) enabling us to characterise the effects of mutations on filament formation (21). This not possible to do for LMM, which forms paracrystals (23). Either homogenous mixtures of either WT or mutant GST LMM molecules (‘homozygous’ conditions) or a 50:50 mixture of WT and mutant LMM molecules (‘heterozygous’ conditions) were mixed together, the resulting filaments were imaged using negative stain EM and the lengths and widths of filaments were measured. Filaments formed under these conditions are not the same as filaments *in vitro*, in which there are also accessory proteins such as titin and MyBPC but they do reveal effects of mutants on the aggregation of LMM into minifilaments.

**Figure 3:**
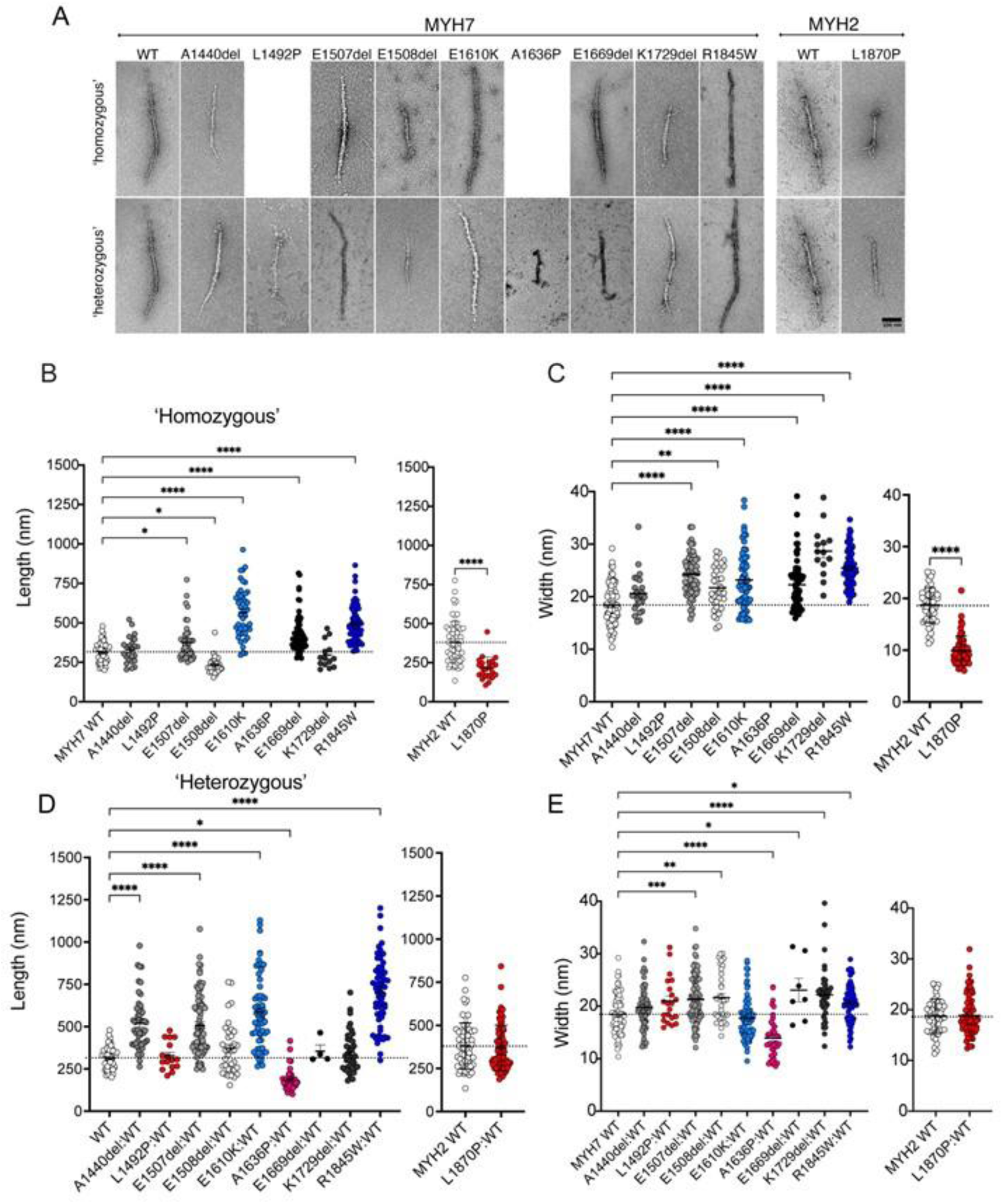
Effects of mutations on aggregation of GST-LMM into minifilaments in vitro. A: shows examples for GST-filaments, negatively stained and imaged using electron microscopy. The top row shows GST-LMM filaments that are effectively ‘homozygous’, as the filaments were generated from a pure population of each type of construct. NB, no images are shown for either of the *MYH7* mutations to proline (L1492P or A1630P) as filaments did not form under these conditions. The lower row shows ‘heterozygous’ GST-LMM filaments, in which the mutant LMM was mixed 50:50 with WT LMM. The images of WT LMM are duplicated for comparison. **B** shows the filament lengths for homozygous populations, and **C** shows the widths. **D** shows filament lengths for heterozygous populations and **E** the widths. Significant differences are as shown. \**p* < 0.05; ** p<0.01; *** p<0.001; *** p<0.0001.

Of the nine MYH7 mutations investigated for homogeneous (homozygous) mixtures of GST-LMM, neither of the proline mutants (L1942P and A1626P) formed minifilaments in vitro (Fig. 3A). When 50:50 mixtures of WT and mutant GST-LMM constructs were mixed (heterozygous), filaments for the two MYH7 proline mutants could now be observed (Fig. 3A). L1492P:WT filaments had similar dimensions to WT filaments, while A1636P:WT filaments were shorter and narrower (Fig. 3D,E). The MYH2 proline mutant (L1870P) did form filaments from homogeneous mixtures, which were shorter and narrower that WT (Fig.3 B,C). When mixed 50:50 with WT, filament dimensions were similar to WT.

The effects of the deletion mutants in MYH7 of GST-LMM filament formation were variable (Fig. 3B-E). In homogeneous mixtures A1440del assembled normally, but in a 50:50 mixture with WT, the filaments were significantly longer. E1507del homogeneous filaments were significantly longer and wider than WT and the same trend was observed in 50:50 mixtures with WT. E1508del homogeneous filaments were significantly shorter and wider than WT, and in 50:50 mixtures, length returned to normal, while width remain increased. E1669del homogeneous filaments were significantly longer and wider than WT and in 50:50 mixtures with WT, length returned to normal while width remained increased. Finally, K1729del homogeneous filaments were significantly wider than WT filaments, as were 50:50 mixtures with WT.

A significant increase in GST-LMM filament length and width was observed for the two missense mutants, E1610K and R1845W, for both homogenous and 50:50 WT:mutant mixtures. Overall, the majority of the mutants tested here have effects on GST LMM aggregation into minifilaments, and these effects are mostly not rescued by mixing in WT GST-LMM.

Finally, we determined the effects of these *MYH7* and *MYH2* mutations on sarcomere incorporation by expressing eGFP-full length myosin heavy chain constructs in differentiated myotubes. In this case, the myosin can co-assemble with endogenous myosin, as well as thick filament accessory proteins to form filaments. Despite effects on filament formation in vitro, all of the mutations we tested were able to incorporate into muscle sarcomeres, with the expected dimensions (Fig. S2). Thus, small disruptions to structure can be accommodated in the cellular environment.

### In the presence of human LMM mutations, myosin heads are disordered

To test whether the myosin mutations have an impact on myosin filament length, myosin head order and myofilament organization *in vivo*, we tested the same mutations in human tissue samples (Table S1). In this case the mutants are heterozygous. Of the 254 muscle fibres (at least 10 myofibres for each individual) analysed, 161 expressed the β/slow myosin heavy chain isoform encoded by *MYH7,* with the remainder expressing the type IIA myosin heavy chain isoform encoded by *MYH2*. Immunofluorescence staining of both patient and control myofibres expressing either β/slow or type IIA myosin heavy chain, showed that muscle sarcomeres were well-preserved, as indicated by regular striated arrays of myosin filaments (Fig. S3A). The length of these filaments (Fig. S3B&C) was consistent with the inter-individual and inter-muscle heterogeneity reported previously (24).

Equatorial reflections related to myosin and actin filaments (1,0 and 1,1, respectively) as well as high quality meridional reflections related to myosin repeats (M3 and M6) were obtained by X-ray diffraction of control and patient muscle bundles in relaxing conditions. Other meridional or layer lines reflections were not analysed as they were difficult to distinguish in the patients. To ensure reliable results and avoid misinterpretation, we pooled all the patients’ data together and compared these to images acquired for controls. This revealed that the intensity ratio (1,1 to 1,0 ratio) was greater in the patients than in the controls, indicating a shift of myosin heads towards the actin filaments. Additionally, the intensity of the M3 layer line, representative of the distance between myosin crowns (typically 14.3nm), was lower in the muscle fibre bundles from patients than in controls, suggesting the heads are more disordered in samples from patients. The intensity of the M6 layer line, which is thought to be related to the helical order of the heads on the thick filament, indicative of myosin filament backbone periodicity/compliance, was preserved (Fig. S4). The spacings of the M3 and M6 meridional layer lines were unchanged between controls and patients (Fig. S4). Altogether, these indicate a specific myosin disorder with heads leaning towards the thin filament.

### In the presence of human LMM mutations, the myosin DRX to SRX ratio is increased

To estimate the ratio of myosin heads in the DRX compared to those in the SRX state, we used Mant-ATP chase experiments (8, 13). This approach relies on a fluorescent ATP named Mant-ATP and on the fact that the rate of ATP consumption by DRX heads is approximately 5 to 10-fold higher than that by SRX heads. Typically, when Mant-ATP is used, the decay of fluorescent intensity over time is best fitted by a double exponential curve (Fig. 4A) allowing the separation of fast and slow phases (indicative of DRX and SRX states, respectively).

**Figure 4.**
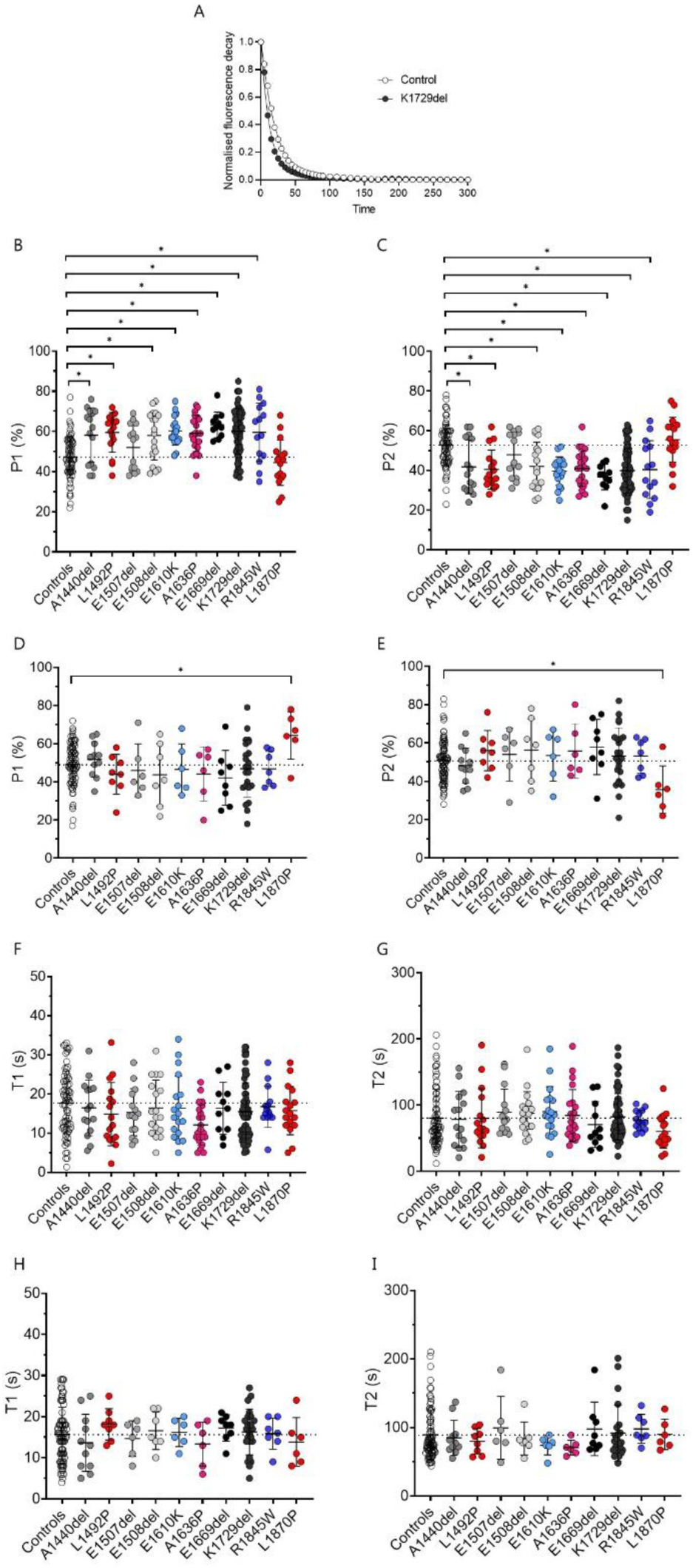
Mant-ATP chase experiments to estimate DRX/SRX ratios. Typical Mant-ATP chase experimental data show exponential decays for muscle fibres (A) isolated from controls (CTL), patients with *MYH7* (MYH7) or with *MYH2* (MYH2) mutations. The proportion of myosin molecules in the disordered-relaxed (P1) and super-relaxed states (P2) as well as their respective ATP turnover lifetimes (T1 and T2) are presented. Data are separated according to the myosin heavy chain expression of individual fibres: either β/slow (graphs B, C, F and G) or type IIA (graphs D, E, H and I). Points for the left panels are individual myofibres. Means and standard deviations also appear on the graphs. \**p* < 0.05.

A total of 456 muscle fibres were tested (more than 20 myofibres for each of the seven controls and for each of the 23 patients, see list in Table S1). Out of these 456 myofibres, 287 expressed the β/slow myosin heavy chain isoform. P1, indicative of DRX, was increased (P2, indicative of SRX, was decreased) in myofibres expressing β/slow myosin heavy chain obtained from patients with *MYH7* mutations, compared with controls, or patient fibres with *MYH2* mutations (Fig. 4B&C). Similarly, P1 was increased (P2 was decreased) in myofibres expressing type IIA myosin heavy chain from patients with *MYH2* mutations compared to controls or patients with *MYH7* mutations (Fig. 4D&E). The ATP turnover lifetime of individual myosin molecules in DRX and SRX states (as assessed by T1 and T2 respectively) was however, not affected in any of the conditions (Fig. 4F-I). Overall, this suggests that, in the presence of heterozygous *MYH7* and *MYH2* mutations, myosin heads are not sequestered properly onto the filament backbone potentially increasing the number of myosin molecules in an ON state ready for crossbridge recruitment (25).

### Modelling suggests that a greater number of myosin motors in the DRX state leads to a small increase in contractility

To predict whether the increased DRX to SRX ratio affects the number of myosin heads binding to actin and the cellular force production, we applied a spatially explicit model of myofilament-level contraction that simulates contractile properties generated by a computational tool, FiberSim 2.1.0 (https://campbell-muscle-lab.github.io/FiberSim/) (26). An increase in myosin heads in the DRX state led to a small increase in isometric force (at maximal and sub-maximal activation levels) and increased Ca^2+^ sensitivity (pCa_50_ = 5.77 compared to pCa_50_ = 5.69) (Fig. 5A&B).

**Figure 5.**
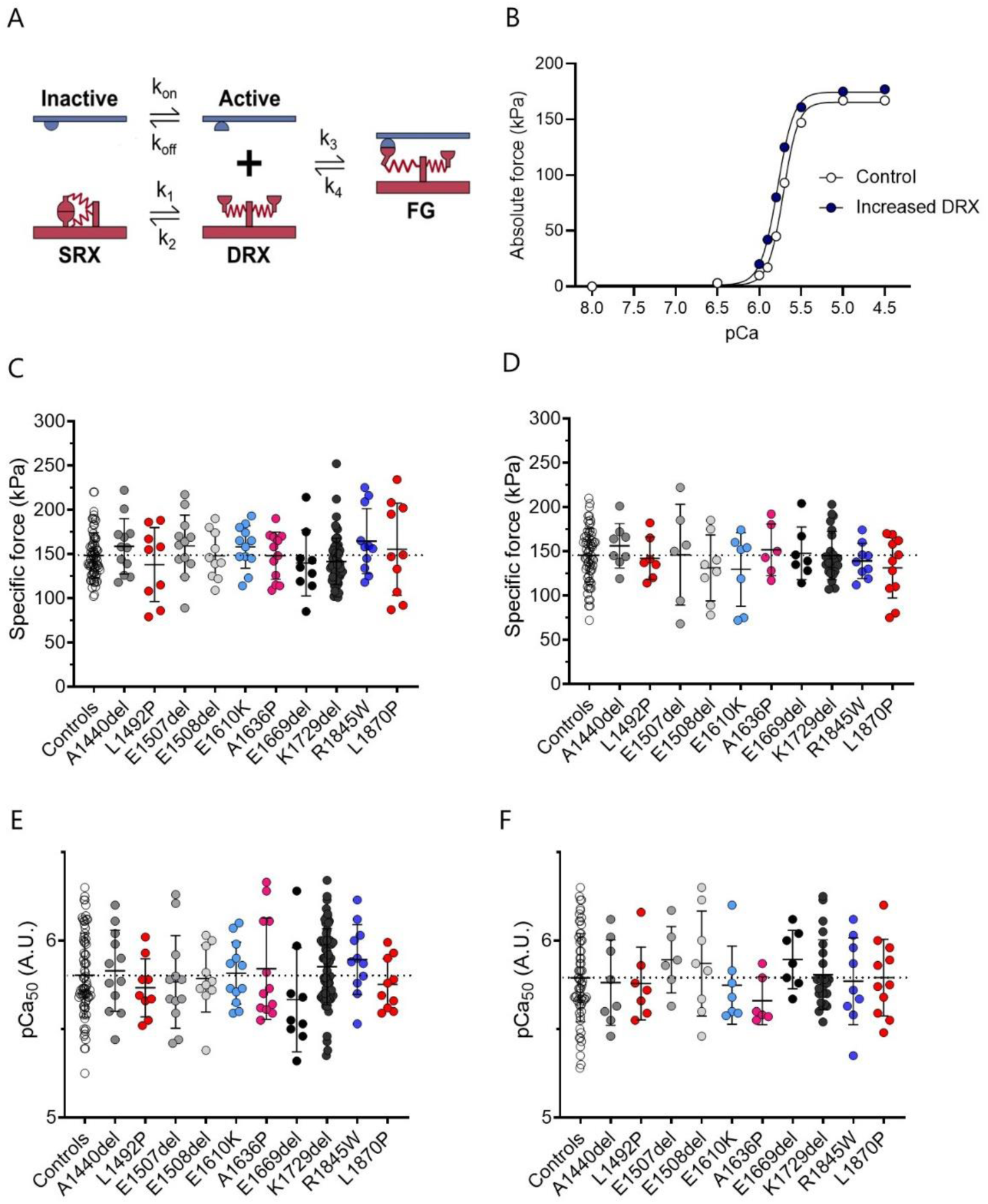
Modelling cellular force production. (A) shows the three-component model used for our simulation from (26) with SRX, DRX and FG (force-generating) states. (B) is the result of the simulations. Experimental specific force and Ca^2+^ sensitivity for muscle fibres isolated from controls (CTL), patients with *MYH7* (MYH7) or with *MYH2* (MYH2) mutations. Data are separated according to the myosin heavy chain expression of individual fibres: either β/slow (graphs C and E) or type IIA (graphs D and F). Points for the left panels are individual myofibres. Means and standard deviations also appear on the graphs.

To complement these computational findings, we measured the contractility of human individual membrane-permeabilised myofibres. A total of 493 myofibres were evaluated (more than 10 myofibres for each of the subjects). Out of these 389 myofibres, 233 were positive for the β/slow myosin heavy chain isoform. However, there were no significant differences in the isometric force produced (Fig. 5C&D) or Ca^2+^ sensitivity (Fig. 5E&F) between myofibres from patients and controls. This likely due to the cell-to-cell variability.

## Discussion

Our findings support the concept that mutations in *MYH7* and *MYH2* that lead to myosinopathy could increase the number of disordered-relaxed heads in relaxed muscle. We found that the majority of the mutations affected secondary structure of the myosin coiled coil and filament formation *in vitro*. Even though the mutant isoforms appeared to incoporated normally in vivo, small effects of the myosin coiled-coil structure have the ability to disrupt packing in the filament. These could induce the destabilisation of shutdown myosin heads, increasing the DRX to SRX ratio, and thus increasing ATP usage in relaxed muscle fibres. Although not tested here, this could also arise through disruption of LMM packing into the filament through its interactions with accessory proteins, such as titin and myosin recently reported in two new CryoEM structures of the C-zone of the thick filament (4, 5). Besides these, we did not observe any significant impact on the muscle fibre force generating capacity. Overall, our results provide important insights into the pathogenesis of skeletal myosinopathies and raise the idea that these particular genetic diseases are under-appreciated and lead to an unexpected increase in ATP usage rather than contractile impairment.

### Increased DRX to SRX ratio as a major pathophysiological mechanism in the presence of LMM mutations

Our observation of altered DRX to SRX ratio is novel in the context of skeletal myosinopathies but has previously been shown to be involved in the pathogenesis of hypertrophic cardiomyopathy arising from *MYH7* mutations. In the latter case, these have been linked to a large number of subtle residue replacements in the myosin motor domains in a region termed the mesa. Mutations in residues in this region interact with the first part of the coiled coil and can directly interfere with the formation of the IHM essential for myosin head sequestration, promoting the cardiac phenotype (27–29), and this has been further explored in a new high resolution structure of shutdown cardiac myosin (30). Here, mutations in the LMM region would need to have a more indirect effect on the stabilisation of IHM heads, through a more general destabilisation of filament packing. An increase in DRX heads, and thus increase in ATP usage, is consistent with clinical findings that demonstrate changes to muscle bioenergetic and metabolic profiles in patients. Specifically, ultrastructural and histological analyses have shown the accumulation, proliferation, and abnormal structure of subsarcolemmal mitochondria (31, 32).

Although more comprehensive studies directly linking myosin metabolic power with mitochondria or whole-body metabolism are needed, an excessive myosin ATP consumption could trigger a metabolic switch away from glycolytic pathways towards a greater reliance on mitochondrial oxidative phosphorylation to meet the increased energy demand. Human resting skeletal muscle metabolic rate is low but accounts for 25% of the obligatory whole-body thermogenesis (33). An increased DRX to SRX ratio by 10-20% (as found here for the *MYH7* and *MYH2* patients) would double skeletal muscle thermogenesis, increasing thermogenesis and the whole-body basal metabolic rate by approximately 16% (8, 13). Regardless of inactivity or limited physical exercise, over a period of a year, the 20% increase in myosin DRX would provoke a mean weight loss of about seven kilograms (13). Altogether, this would support clinical observations reporting that patients diagnosed with skeletal myosinopathies who are lean or underweight, despite being inactive and sedentary.

### Dysregulated coiled-coil structure and filament packing as a potential cause of the increased DRX to SRX ratio

In contrast to *MYH7*-driven hypertrophic cardiomyopathy (27–29), here, the reduced myosin head sequestration in patients cannot be attributed to a direct effect of the mutations on the interacting heads motif through the myosin mesa but rather to other processes involving the LMM region. LMM is highly conserved among vertebrates emphasizing the importance of its amino acid arrangement for the formation of the heptad repeats (i.e., *a*-*g*) within the coiled-coil and for myosin filament packing (34). Interestingly, here, most of the subtle *MYH7* and *MYH2* mutations affect residues in *b*, *c* and *f* positions of the charged exterior portion of the coiled coil (34). Of the mutations we have studied here, the deletion mutations in E1507, E1508, E1669 and K1729 occur in regions of LMM which are strongly negatively charged (Fig. S7). Deletion of these residues would be expected to decrease the overall negative charge of this region in addition to disruption of the coiled-coil structure, and both of these factors could contribute to altered filament packing in thick filaments. E1610 and R1845 are both in regions that are of strong positive charge and likewise have the potential to disrupt myosin packing into thick filaments through this change to the overall positive charge.

In striated muscle myosins, the interacting heads motif is thought to mainly depend on the interaction of the two heads with each other, and with the first part of the coiled coil (S2). However, the proportion of myosin heads in the SRX state appears to strongly depend on the molecular state of the myosin, being low in single molecules and much higher in molecules organized into filaments (reviewed in (27)) where myosin molecules can interact with tails in the filament backbone, as well as with thick filament proteins such as myosin-binding protein C and titin. This raises the possibility that interactions with LMM may also play a role in stabilizing the SRX state in the filament. In isolated, shutdown, smooth muscle myosin, the distal coiled coil region (LMM) interacts with the so-called blocked head of the interacting heads motif, with key interactions between LMM and the SH3-like fold at the N-terminus of the motor domain, and with the converter domain (35, 36). A similar interaction between LMM and the IHMs in cardiac and skeletal muscle filaments could help to keep the myosin heads in the IHM state, by inhibiting the movement of the converter domain of the blocked head (27). Our results indicate that while the mutations likely affect the local structure of the coiled coil, they do not have significant effects on sarcomeric incorporation in vivo. However, they could affect the ability of the myosin to adopt an SRX state through a subtle disruption of myosin filament packing.

## Conclusion

Taken together, our findings demonstrate a clear pathogenic effect of *MYH7* and *MYH2* mutations (associated with congenital myopathies) on myosin coiled-coil structure. Additionally, our results highlight that, in resting human myofibres from patients, the myosin-stabilizing conformational state is altered increasing the DRX to SRX ratio and basal ATP demand. All these do not significantly impact the cellular force producing capacity. Nevertheless, they give new valuable insights into skeletal myosinopathies, the unexplained odd appearance of energetic proteins (e.g., mitochondria) and the potential benefits of myosin-linked drugs targeting its ATPase activity.

## Materials and Methods

### Generation of LMM mutant constructs

The cDNA used in these experiments encodes sequence from the human *MYH7* (P12883, Uniprot) and MYH2 (Q9UKX2, Uniprot). The cDNA for *MYH7*, and the *MYH7* LMM (residues 1280–1936) constructs were generated as described previously (21). The LMM coding region was subcloned into pGEX-6P-1 (GE Lifesciences) as previously described (21), which adds a GST sequence and a PreScission protease site at the N-terminus. The desired mutations for *MYH7* were introduced using the Quick ChangeXL II Site Directed Mutagenesis Kit (Agilent) according to the manufacturers’ instructions. Each construct was sequenced to verify that the desired mutations had been introduced and to confirm that no other mutations had been introduced. The cDNA for full length *MYH2*, together with a mutated form (L1870P) was obtained from Genscript. The cDNA for *MYH2* LMM region (residues 1223-1941) was subcloned into the pGEX-6P-1 (GE Lifesciences) for expression and purification. All LMM constructs were designed to start and finish on an amino acid in the d position of the heptad repeat.

### Expression and purification of LMM

The WT and mutant constructs were transformed into *E. coli* Rosetta 2 (Novagen), a single colony was inoculated into a 5-ml starter culture and grown overnight before being added to 500 ml Auto-Induction media (Formulation (g/l) - 12g Tryptone, 24g Yeast extract, 0.15g MgSO_4_, 3.3g (NH4)_2_SO_4_, 6.5g KH_2_PO_4_, 7.1g Na_2_PO_4_, 0.5g Glucose, 2g Alpha Lactose pH 7.0) and grown at 20°C for 18 hours. Cells were harvested and the pellets stored at −80 °C. For protein purification, the pellets were resuspended in lysis buffer [PBS, 350 mM NaCl, 1 mM DTT, 1 mM EDTA, 200 μg/ml lysozyme, 0.1% Triton X-100, protease cocktail inhibitor tablet (Roche; pH 7.5)] for 30 min at room temperature on a roller. The lysates were sonicated on ice (6 cycles of 10 s on, 10 s off) and the cell debris was pelleted by centrifugation (30000g, 30 min). Proteins were purified by GST-tag affinity chromatography using Glutathione Sepharose 4B (GE Lifesciences). First, cleared lysates were added to the equilibrated 1.5-2 ml glutathione Sepharose 4B and washed with 100ml of wash buffer (PBS, 350 mM NaCl, 1 mM DTT, 1 mM EDTA, pH 7.5). For CD experiments, the LMM was liberated from the column matrix by overnight incubation with PreScission Protease in cleavage buffer [50 mM Tris–HCl, 150 mM NaCl, 1 mM EDTA, 1 mM DTT (pH 7.5)] at 4 °C. The constructs were then concentrated with a vivaspin 6 and dialysed against CD buffer (500 mM NaCl, 10 mM phosphate buffer (pH 7.5), 1 mM DTT). For EM studies, the GST tag was not cleaved away from LMM constructs, and the intact GST–LMM was eluted from the column by using 20 mM reduced glutathione. To assess the purity of the protein, fractions were collected, the protein content and purity were assessed by SDS-PAGE, and protein concentration was measured using the BCA Assay (Sigma).

### Circular Dichroism (CD) measurements

CD spectra for LMM constructs from which GST had been removed were measured at a temperature of 10°C at wavelengths from 260 to 190 nm using an APP Chirascan CD spectropolarimeter. The CD buffer contained 500 mM NaCl, 1 mM DTT and 10 mM phosphate buffer (pH 7.5). The concentrations for all LMM constructs were 150–250 μg/ml. Scans at 10 °C were repeated twice, and a minimum of three experiments were performed. To calculate any significant difference, t tests of the 222-nm MRE value were used. A fully (100%) helical construct was considered to have an MRE value at 222 nm of 36,000 (100%) (Greenfield & Fasman, 1969). This value was used to estimate the % helical content of the constructs. Thermal melting measurements were taken at 1°C increments from 10 to 80°C, with 1°C/min heating rate, in the same buffer.

### Electron Microscopy (EM)

GST–LMM filaments were generated by rapidly diluting GST-LMM into a low salt solution (150 mM NaCl, 10 mM MOPS (pH 7.2), 1 mM EDTA). The mixture was then left on ice for 2 min to allow for filament formation. The final concentration of GST–LMM was 0.3–0.5 μM. 5 µl was then loaded onto a carbon-coated copper grid, previously irradiated using a PELCO easiGlow Glow Discharge Cleaning system (TedPella). The sample was allowed to adhere for 10 seconds before the sample was flicked away and stained with 1% uranyl acetate. The grids were imaged using an FEI Tecnai TF20 microscope at 120 kV and micrographs were recorded using a 4k × 4k Gatan CCD camera (pixel size of 0.351 nm). Measurements of filament lengths were performed using ImageJ (FiJi) software (37). Graphs and analysis were generated using Prism (Graph Pad) software.

### Molecular Dynamic Simulations

Simulations were performed as previously described (21). Simulations using explicit solvent were performed using the CHARMM-36 force field parameters (38) with TIP3P water on fragments that were 13 heptads (91 residues long). A starting structure for the composite model of residues 1526–1689 within the LMM region of human β-MHC (39) was generously supplied by Professors Ivan Rayment and Qiang Cui. MD simulations were also performed on shorter segments of this same part of the β-MHC coiled coil for which experimental atomic structures are available (40). For these, the N-terminal globular Xrcc4 moieties were removed using Chimera (41). Specifically, residues 1562–1608 were taken from 5CJ4 (chain A: residues 1562–1615 and chain B: residues 1562–1608, (40): residues 1590–1657 were taken from 5CHX (40), and residues 1631–1689 were taken from 5CJ0 (39). Methylated lysine side chains (modified to facilitate crystallization in 5CJ4 and 5CHX) were converted back to unmodified lysine. In all cases, N-termini were capped with acetyl groups and C-termini were capped with N-methylamide for both chains. Atomic structures equivalent to the WT composite models were also made for all mutants. To generate the substitution mutants, the most probable rotamer for each chain was substituted using the ‘Structure Edit’ function within Chimera (41).

For skeletal myosin, the coiled-coil tail model (7KOG) from the *Lethocerus indicus* thick filament cryo-EM reconstruction (42) was used as a starting model. The skeletal myosin sequence was threaded onto the 7KOG model using Modeller 9.18 (43). A 13 heptad segment (residues 1825 – 1915) was extracted to yield a WT model. The L1870P mutant was generated by mutating L1870 to a proline using the “Structure Edit’ function within Chimera as before.

Generation of a starting structure for the deletion mutants was performed as described previously (21). Producing a model for deletion mutants was more challenging than for the substitution mutants, since simply deleting a residue from the WT model and rebonding the chains across the gap produces a break in the coiled-coil structure that would not occur during formation of the mutant coiled coil in vivo. We therefore built a non-canonical coiled coil ab initio for each deletion mutant, along with a corresponding WT equivalent model using the program BEAMMOTIFCC (22). These models used noncanonical structures for the skip residue (where applicable) and for the deletion site. Briefly, the initial WT model was built using the following values: 3.617 residues per turn of helix, an axial translation per residue of 1.495 Å, and a relative rotation of the two helical strands of 210°. The major helical radius used was 4.9 Å, which was the average major helical radius calculated along the coiled coil from simulations of the composite PDB model. The smoothing parameter b was varied to find a value that resulted in simulation average properties that best matched those from simulating the composite PDB model (b = 0.03 was the optimum). The structure is defined in the program by a pattern of “motifs” that link equivalent residues in the sequence; the motif for a canonical coiled coil being 7-amino-acid residues. To accommodate the skip residues E1582 and G1807, a 29-residue motif replaced four 7-residue motifs between F1565 and V1594, as well as L1796 and E1825, respectively. We validated the method by comparing these 91-residue WT models with the composite model, where applicable.

Starting simulations from our ab initio model gave rise to average properties that all matched very well those found for simulations initiated from the composite model, showing that the method copes well with skip residues. We then used the same approach to build deletion coiled-coil mutants with a smooth all-helix structure from which to initiate simulations. To build a non-canonical deletion model with a continuous helical structure, in addition to the skip motif modification (where applicable), a 27-residue motif replaced four canonical 7-residues motifs. The BEAMMOTIFCC program was modified in order to implement the 27-residue motif, which is defined to contain 8 helical turns (like in the WT four 7-residue motifs that it replaced) to ensure a left-handed coiled coil (refer to the N value in (Offer et al., 2002) for details). BEAMMOTIFCC provides a backbone structure for the coiled-coil model. Side-chain atoms were added to the model using SCWRL4 (44). N-termini were again capped with acetyl groups and C-termini were capped with N-methylamide. Simulations were run as for the composite model with an additional minimization and equilibration run with restrained backbone atoms to allow for side-chain only equilibration (10,000-step minimization, 0–300 K heating protocol and 100,000 step pre-equilibration performed using AMBER) prior to the all atom minimization.

CHARMM-GUI (45) was used to add a 1.5-nm surround of water molecules and Na^+^ and Cl^−^ ions were then added to neutralize the construct and give a NaCl concentration of ∼ 150 mM. A 10,000-step minimization, 0–300 K heating protocol and short pre-equilibration run (100,000 steps) were performed using AMBER. Data are taken from simulation runs lasting at least 1 µs across at least 3 separate runs initiated using a random number seed in AMBER (46) at 300 K. The timestep used was 4 fs and trajectory frames were recorded every 25000 steps. The first 10 ns of each run was removed prior to analysis to avoid starting structure bias. Simulation trajectories were analyzed using CPPTRAJ (47). The helicity of both chains was calculated using backbone dihedral angles and the method previously described (38). The mean ± SD distances between helices (Dcom) values were calculated using a moving window to include the positions of seven Cα (3 N-terminal and 3 C-terminal to the marked residue in each chain). Similarly, average heptad lengths along each helix were calculated using a moving window average of Cα positions in seven (i) residues and their (i + 7) residue partners in the sequence. The inter-heptad angles were measured between Cα atoms in residues at sequence positions (i – 7), i, and (i + 7).

### Expression of eGFP–β-MHC in cardiomyocytes and myotubes

C2C12 myoblasts, purchased from Public Health England culture collections, were used to generate skeletal muscle myotubes in culture. They were maintained in Dulbecco’s minimum essential medium (DMEM), with high glucose and Glutamax, supplemented with 20% FCS and 1% penicillin and streptomycin (diluted from stocks containing 10,000 U/ml) and used at low passage (<10). The cells were differentiated in DMEM supplemented with 2% horse serum and 1% penicillin and streptomycin. The adenovirus expression vector, pDC315, was used to express the full-length heavy chain of MYH7 (P12883, Uniprot) and full-length heavy chain of MYH2 (Uniprot: Q9UKX2). In both constructs, eGFP was fused to the N-terminus. Virus was generated, purified using Vivapure AdenoPACK 100 (Sartorius), and titred as described previously (21). Average titres were 2×10^8^ pfu/ml.

C2C12 myoblasts were seeded at the same cell density (1 × 10^5^ cells/ml) on coverslips coated with laminin-1 (Sigma). 24 hours after seeding, cells were infected with 5µl of eGFP-MHC adenovirus together with 5µl of mCherry-UNC45b Co-Chaperone adenovirus (equivalent to MOIs of ∼10). The addition of this co-chaperone improved sarcomere incorporation of eGFP-MYH2 and MYH7. After 24-h, the growth medium was exchanged for differentiation medium and the cells were incubated at 37°C for 5 days to allow for differentiation into skeletal muscle myotubes. The cells were then fixed with 2% PFA for 20 mins at room temperature as described (21).

### Immunostaining and microscopy

Cells were stained with DAPI (Molecular Probes) to visualize the nucleus. Images of eGFP-myosin expressing cells were obtained using a Olympus BX51 microscope and a x100 objective (N.A. 1.4) to perform an analysis of sarcomere organization. The same camera settings were used to capture all the images. Additional images were obtained using a Zeiss LSM 880 confocal with Fast Airyscan microscope, using a x40 objective (N.A. 1.4).

The images of eGFP–MHC in skeletal myotubes were analysed using ImageJ to assess sarcomeric incorporation. Lines of a fixed width were drawn along 4–5 sarcomeres in the same myofibril, and the intensity profile along the line was measured using the plot profile function. Intensity profiles were imported into an Excel spreadsheet, manually aligned to the minimum value in the centre of the sarcomere for each sarcomeric measurement and used to generate an average plot for all the sarcomeres in that myofibril. The average values for each myofibril were then combined and averaged across myofibrils, again aligning on the minimum value. Values for a minimum of 30 sarcomeres were measured for WT and each mutant and used to calculate the overall mean values and generate the profile plots.

### Human subjects

Muscle biopsy specimens were obtained from patients diagnosed with skeletal myosinopathies and with either *MYH7* or *MYH2* mutations; and from age-matched controls with no history of neuromuscular diseases. Details of all the patients are given in Table S1. All samples were flash-frozen and stored at - 80°C until analysed.

### Solutions

Relaxing and activating solutions contained 4 mM Mg-ATP, 1 mM free Mg^2+^, 20 mM imidazole, 7 mM EGTA, 14.5 mM creatine phosphate and KCl to adjust the ionic strength to 180 mM and pH to 7. Additionally, the concentrations of free Ca^2+^ ranged 10^-9^ M (relaxing solution, pCa 9) to 10^-4.5^ M (maximum activating solution, pCa 4.5). The rigor buffer for Mant-ATP chase experiments contained 120 mM K acetate, 5 mM Mg acetate, 2.5 mM K_2_HPO_4_, 50 mM MOPS, 2 mM DTT with a pH of 6.8 (48).

### Muscle preparation and fibre permeabilisation

Cryopreserved muscle samples were immersed in a membrane-permeabilising solution (relaxing solution containing glycerol; 50:50 v/v) for 24 hours at -20°C, after which they were transferred to 4°C and bundles of approximately 50 muscle fibres were dissected free and then tied with surgical silk to glass capillary tubes at slightly stretched lengths. These bundles were kept in the membrane-permeabilising solution at 4°C for an additional 24 hours (to allow a proper skinning process). After these steps, bundles were stored at -20°C for use up to one week (48).

### **X-** ray diffraction recordings and analyses

On the day of the experiments, bundles were placed in a plastic dish containing the relaxing solution. They were then transferred to the specimen chamber, filled with the relaxing buffer. The ends of these thin muscle bundles were then clamped at a resting sarcomere length (≈2.20 µm). Subsequently, X-ray diffraction patterns were recorded at 15°C using a CMOS camera (Model C11440-22CU, Hamamatsu Photonics, Japan, 2048 x 2048 pixels) in combination with a 4-inch image intensifier (Model V7739PMOD, Hamamatsu Photonics, Japan). The exposure time was 500 ms. The X-ray wavelength was 0.10 nm and the specimen-to-detector distance was 2.14 m. For each preparation, approximately 20-30 diffraction patterns were recorded at the BL40XU beamline of SPring-8 and were analysed as described previously (49). To minimize radiation damage, the exposure time was kept low and the specimen chamber was moved by 100.00 μm after each exposure. Following X-ray recordings, background scattering was subtracted, and the equatorial reflections as well as the major myosin meridional reflection intensities/spacing were determined as described elsewhere previously (49, 50).

### Mant-ATP chase experiments

On the day of the experiments, bundles were transferred to the relaxing solution and single myofibres were isolated. Their ends were individually clamped to half-split copper meshes designed for electron microscopy (SPI G100 2010C-XA, width, 3.00 mm), which had been glued to glass slides (Academy, 26.00 x 76.00 mm, thickness 1.00-1.20 mm). Cover slips were then attached to the top (using double-sided tape) to create flow chambers (Menzel-Gläser, 22.00 x 22.00 mm, thickness 0.13-0.16 mm) (48). Subsequently, at 25°C, for each muscle fibre, the sarcomere length was checked using the brightfield mode of a Zeiss Axio Scope A1 microscope. One sarcomere length (≈2.20 µm) was used in the present study and further subjected to the Mant-ATP chase protocol. Similar to previous studies (29, 48, 51) each fibre was first incubated for 5 minutes with a rigor buffer. A solution containing the rigor buffer with 250 μM Mant-ATP was then flushed and kept in the chamber for 5 minutes. At the end of this step, another solution made of the rigor buffer with 4.00 mM ATP was added with simultaneous acquisition of the Mant-ATP chase. For fluorescence acquisition, a Zeiss Axio Scope A1 microscope was used with a Plan-Apochromat 20x/0.8 objective and a Zeiss AxioCam ICm 1 camera. Frames were acquired every 5 seconds with a 20 ms acquisition/exposure time using a DAPI filter set, images were collected for 5 minutes (tests were run prior to starting the current study where images were collected for 15 min instead of 5 min – these tests did not reveal any significant difference in the parameters calculated). Three regions of each individual myofibre were sampled for fluorescence decay using the ROI manager in ImageJ as previously published (29, 48, 51). The mean background fluorescence intensity was subtracted from the average of the fibre fluorescence intensity (for each image taken). Each time point was then normalized by the fluorescence intensity of the final Mant-ATP image before washout (T = 0). These data were then fit to an unconstrained double exponential decay using SigmaPlot 14.0 (Systat Software Inc):

Where P1 is the amplitude of the initial rapid decay approximating the ‘disordered-relaxed state’ (DRX) with T1 as the time constant for this decay. P2 is the slower second decay approximating the proportion of myosin heads in the ‘super-relaxed state’ (SRX) with its associated time constant T2 (29, 48, 51).

### Single muscle fibre contractility

As for Mant-ATP experiments, individual myofibres were dissected in the relaxing solution. They were then individually attached between connectors leading to a force transducer (model 400A; Aurora Scientific) and a lever arm system (model 308B; Aurora Scientific). Sarcomere length was set to ≈2.50 µm and the temperature to 15°C (48). As the baths had glass bottom and right angle prisms, fibre cross-sectional area (CSA) could be estimated from the width and depth, assuming an elliptical circumference. To determine the maximal and submaximal isometric force generating capacity, myofibres were sequentially bathed in activating buffers with increasing [Ca^2+^] termed pCa. Specific force corresponded to absolute force normalized to myofibre cross-sectional area. Submaximal force values were normalised to the maximal force and the obtained force-pCa curve were fit to the Hill equation (four-parameters). Ca^2+^ sensitivity was then obtained and corresponded to the pCa at which 50% of maximal force is reached. *n*H was also calculated and represented the steepness of the force-pCa curve indicative of the degree of actin-myosin cross-bridge cooperativity (48).

### Immunofluorescence staining and confocal imaging

To avoid any potential misinterpretation due to the type of myosin heavy chain (*MYH7*: β-cardiac/skeletal slow myosin heavy chain or *MYH2*: skeletal fast myosin heavy chain 2A), for all the above experiments in individually isolated muscle fibres, we assessed the sub-type right after the Mant-ATP chase experiments or contractile measurements using immunofluorescence staining and confocal imaging as previously described (48). Briefly, the ‘used’ individual myofibres were individually mounted once again as for the Mant-ATP chase experiments and incubated with A4.951 (1:25, Mouse IgG1, DSHB) or SC71-S (1:25, Mouse IgG1, DSHB) followed by Alexa Fluor 488 or 647 (1:500, Goat anti-mouse, ThermoFisher Scientific). Images were acquired using a confocal microscope (Zeiss Axiovert 200, objectives ×20, ×40, and ×100) equipped with a CARV II confocal imager (BD Biosciences) (48). Distributed deconvolution (DDecon) was then applied from the acquired images with a specific plugin for ImageJ (National Institutes of Health, Bethesda, MD) (52). Note that DDecon is a super-resolution light microscopy technique that allows the computation of filament lengths with a precision of 10.00–20.00 nm (52). All line scans were background-corrected. Distances (and myosin filament lengths) were finally calculated by converting pixel sizes into micrometer using the magnification factor for each image (52).

### Myofilament model and simulation

The FiberSim 2.1.0 software has been thoroughly described (26). Briefly, FiberSim tracks the position and status of actin and myosin molecules within a network of compliant thick and thin filaments. For all the simulations, the half-sarcomere lattice was composed of 100 thick filaments and 200 thin filaments. These filaments were arranged in a hexagonal lattice to mimic the architecture of human myofibres. Filaments located at the edge of the lattice were “mirrored” on the opposite side to minimize edge effects (53). Each thin filament was composed of two actin strands. Each strand contained 27 regulatory units, and each regulatory unit contained 7 binding sites. Each thick filament was composed of 54 myosin crowns with each crown consisting of 3 pairs of myosin dimers. Each binding site on actin could be in an inactive (unavailable for myosin binding) or active (available for myosin binding) state. All 7 binding sites from a regulatory unit switched simultaneously between those two states, depending on the Ca^2+^ concentration and on the transition rate constants k_on_ and k_off_ which we set. A cooperative mechanism was also implemented such that the transition probability for a regulatory unit was influenced by the states of its neighbours. Finally, a regulatory unit was prevented from deactivating if one or more myosin head(s) were bound.

Although this model only has two explicit thin filament states, it mimics an important feature of the three state thin filament model described by McKillop and Geeves (54) in that bound myosin heads inhibited relaxation. Heads in SRX switched to DRX at a rate k_1_ that is assumed to be force-dependent (26). DRX heads could then attach to available binding sites on actin. The attachment and detachment rates depended on x, where x is the distance to the binding site measured parallel to the filaments. The rate functions are provided in Table S2. The detachment rate function (k_4_) was updated for the present study so that it had an exponential strain-dependence similar to that measured for single myosin heads via optical trapping (55). Model parameters were chosen to reproduce physiological values for: maximal isometric force (≈ 150-200 kPa at a sarcomere length of 2.2 µm), passive force (≈ 1-2% of the maximal isometric force) and Ca^2+^ sensitivity (pCa_50_ ≈ 5.7).

### Statistics

Data are presented as means ± standard deviations. Graphs were prepared and analysed in Prism 9.0 (GraphPad). Statistical significance was set to *p* < 0.05. Control data points were pooled, since no significant differences were observed amongst healthy control human subjects. Owing to the different origins of disease (mutation and gene affected), patient data points were not pooled. One-way ANOVA with Tukey post-correction was used to compare each patient against the pooled controls; in addition, a random effect algorithm was incorporated into the model, to account for any potential inter-individual differences that might exist amongst the control cohort (56).

### Study approval

All tissue was consented, stored and used in accordance with the Human Tissue Act, UK, under local ethical approval (REC 13/NE/0373).

### Data availability

All the raw data of this manuscript are shown in the figures but if needed, these can be made available upon request.

## Acknowledgments

We thank the Nikon Imaging Centre at King’s College London for the provision of equipment for, and assistance with confocal imaging. The X-ray experiments were performed under approval of the SPring-8 Proposal Review Committee (2020A1050 and 2021B1085).

This work was generously funded by the Medical Research Council UK to M.P. and J.O. (MR/S023593/1), Muscular Dystrophy UK (17GRO-PS48-0077) to J.O.; National Institute of Health (HL148785) to K.S.C and Estonian Research Council (PUT355, PSG774 and PRG471) to S.P. and K.O. The confocal microscope at Leeds was funded by the Wellcome Trust (104918/Z/14/Z).

## Authors’ contributions

GC, SK, KSC, MP and JO designed the work; GC, AH, SK, HFD, FM, ADA, MM, PVDB, CF, NBR, EM, EP, JJV, AO, SP, KO, MGP, MR, EZ, WDR, KSC, HI, MP and JO performed data collection; GC, AH, SK, HFD, KSC, HI, MP and JO ran data analysis and interpretation; GC, MP and JO drafted the article; GC, AH, SK, HFD, FM, ADA, MM, PVDB, CF, NBR, EM, EP, JJV, AO, SP, KO, MGP, MR, EZ, WDR, KSC, HI, MP and JO critically revised the article; GC, AH, SK, HFD, FM, ADA, MM, PVDB, CF, NBR, EM, EP, JJV, AO, SP, KO, MGP, MR, EZ, WDR, KSC, HI, MP and JO approved the final version.

