## Supplementary File for "Human light meromyosin mutations linked to skeletal myopathies disrupt the coiled coil structure and myosin head sequestration"

**Table S1**
Patient and control muscle biopsy samples used.

| Age (years) | Gender (M/F) | Mutation | Amino acid change | Heptad position | Disease | Source |
|---|---|---|---|---|---|---|
| | | | *MYH7* | | | |
| 15 | F | c.4319delTGC | p.A1440del | *b* | Laing Distal Myopathy | London, UK |
| 16 | F | c.4475T>C | p.L1492P | *e* | Laing Distal Myopathy | Rome, Italy |
| 26 | M | c.4520-15_4520-9del | p.E1507del | *f* | Congenital Myopathy | Paris, France |
| 68 | F | c.4522_4524del | p.E1508del | *g* | Laing Distal Myopathy | Brussels, Belgium |
| 44 | F | c.4828G>A | p.E1610K | *c* | Congenital Myopathy | Paris, France |
| 48 | F | c.4906G>C | p.A1636P | *g* | Laing Distal Myopathy | Valencia, Spain |
| 49 | F | c. c.5005_5007delGAG | p.E1669del | *f* | Laing Distal Myopathy | Gothenburg, Sweden |
| 31 | F | c.5292_5294delAGA | p.K1729del | *c* | Laing Distal Myopathy | Valencia, Spain |
| 42 | M | c.5292_5294delAGA | p.K1729del | *c* | Laing Distal Myopathy | Valencia, Spain |
| 48 | F | c.5292_5294delAGA | p.K1729del | *c* | Laing Distal Myopathy | Valencia, Spain |
| 57 | M | c.5292_5294delAGA | p.K1729del | *c* | Laing Distal Myopathy | Valencia, Spain |
| 60 | F | c.5292_5294delAGA | p.K1729del | *c* | Laing Distal Myopathy | Valencia, Spain |
| 60 | F | c.5292_5294delAGA | p.K1729del | *c* | Laing Distal Myopathy | Valencia, Spain |
| 62 | F | c.5292_5294delAGA | p.K1729del | *c* | Laing Distal Myopathy | Valencia, Spain |
| 65 | F | c.5292_5294delAGA | p.K1729del | *c* | Laing Distal Myopathy | Valencia, Spain |
| 74 | M | c.5292_5294delAGA | p.K1729del | *c* | Laing Distal Myopathy | Valencia, Spain |
| 57 | M | c.5533C>T | p.R1845W | *f* | Myosin Storage Myopathy | Valencia, Spain |
| | | | *MYH2* | | | |
| 1 | M | c.5609T>C | p.L1870P | *d* | Congenital Fibre Type Disproportion | Tartu, Estonia |
| | | | *Controls* | | | |
| 10 | M | - | - | | - | Sao Paulo, Brazil |
| 20 | M | - | - | | - | London, UK |
| 20 | F | - | - | | - | Sao Paulo, Brazil |
| 25 | F | - | - | | - | London, UK |
| 32 | M | - | - | | - | Copenhagen, DK |
| 54 | F | - | - | | - | Copenhagen, DK |
| 71 | F | - | - | | - | Copenhagen, DK |

**Table S2**

Parameters used for the simulation.

| Actin kinetics | Parameters |
|---|---|
| $k_{activate} = [Ca^{2+}] \cdot k_{on} \cdot (1+ n \cdot \gamma_{coop})$<br>$k_{deactivate} = k_{off} \cdot (1 + [2 - n] \cdot \gamma_{coop})$ | $k_{on} = 2.\ 10^7\ M \cdot s^{-1}$<br>$\gamma_{coop} = 10$<br>n = 0,1 or 2 (# of active neighboring RUs)<br>$k_{off} = 100\ s^{-1}$ |
| **Myosin kinetics** | |
| $k_1 = k_{1,0} + k_{1,f}$ . node force | $k_{1,0} = \begin{cases} 45\ s^{-1} \text{ for control} \\ 100\ s^{-1} \text{ for mutation} \end{cases}$<br>$k_{1,f} = 100\ nN\ s^{-1}$ |
| $k_2$ = constant | $k_2 = 100\ s^{-1}$ |
| $k_3 = k_{3,0} \exp(-\frac{k_{cb} x^2}{2\ k_B T})$ | $k_{3,0} = 25\ s^{-1}$<br>$k_{cb} = 10^{-3}\ nN.nm^{-1}$<br>T = 310 K |
| $k_4 = \begin{cases} k_{4,0} \exp(-k_{4,1}(x + x_{ps})) \text{ if } x < x_{wall} \\ 1000 \text{ if } x > x_{wall} \end{cases}$ | $k_{4,0} = 150\ s^{-1}$<br>$k_{4,1} = 0.25\ nm^{-1}$<br>$x_{ps} = 5$ nm<br>$x_{wall} = 8$ nm |

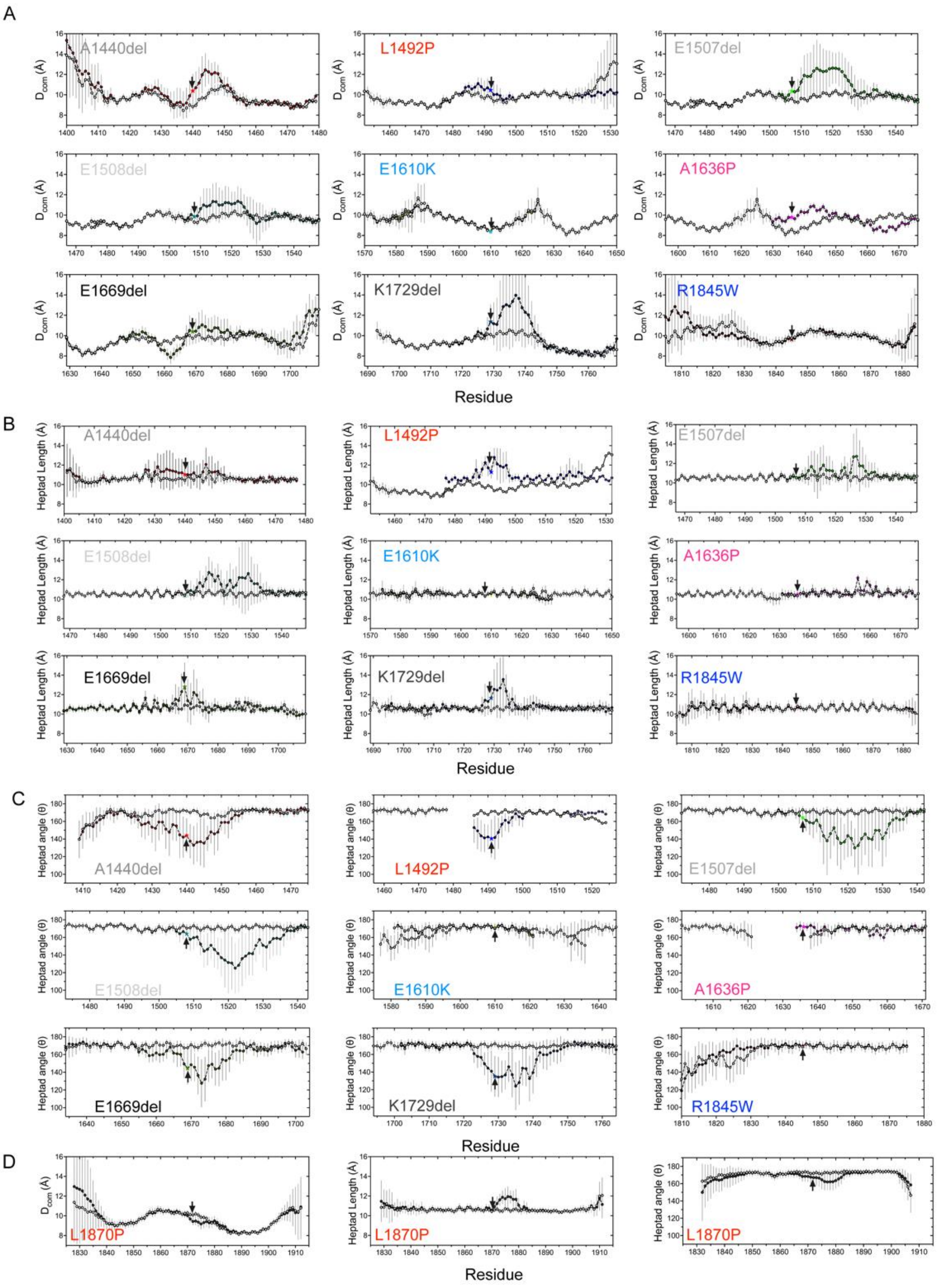


**Figure S1: Results from MD simulation of WT and mutant coiled coil models.** **A.** The distance between the helices ($D_{com}$) for each MYH7 mutant (indicated by star) as a function of residue position compared to WT (white). **B.** Heptad length, the distance between Cα atoms in residues at positions *i* and *i* + 7, averaged across both chains of the coiled-coil and **C:** Inter-heptad angle, the angle between the lines linking Cα atoms in residues at positions (*i* – 7), (*i*) and (*i*, *i* + 7), averaged across both chains of the coiled-coil. **D.** The distance between the helices ($D_{com}$) (left), Heptad Length (centre), and Heptad angle (right), for MYH2 mutant, L1871P (indicated by star) as a function of residue position compared to WT (white). Error bars represent the SD during the simulation.

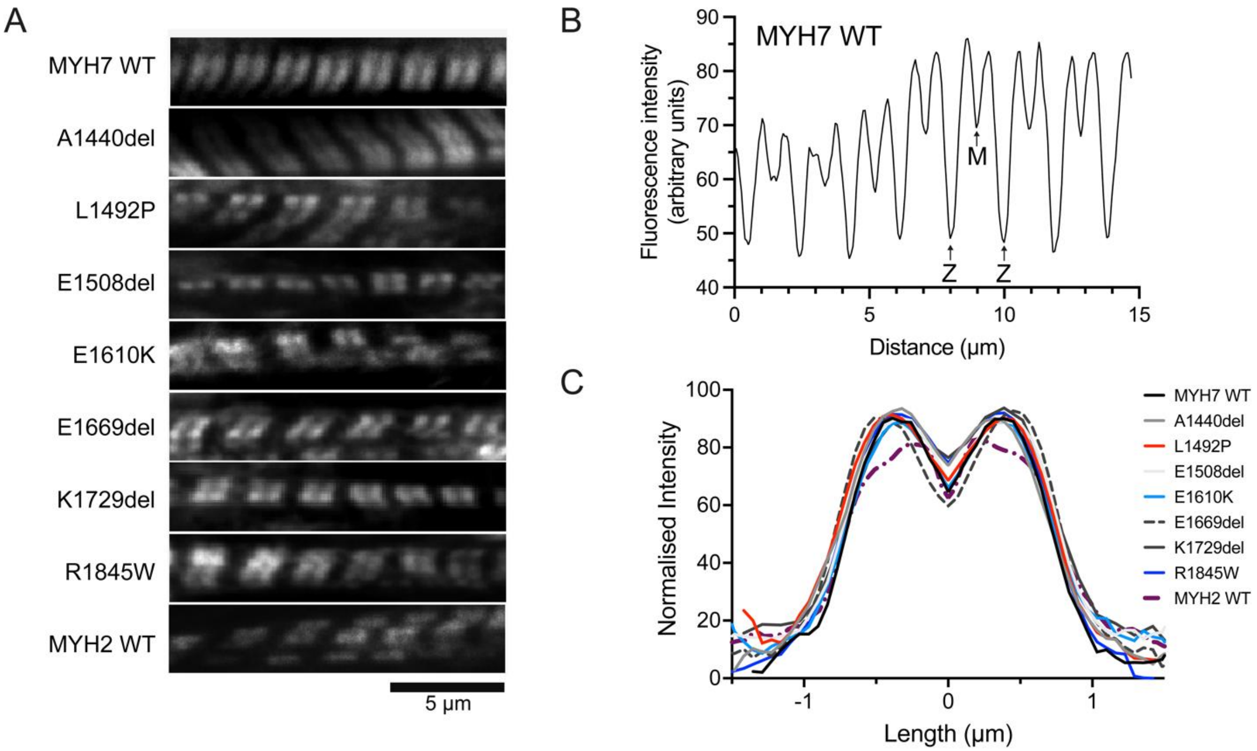


**Figure S2: Mutant isoforms of eGFP–MHC do not affect incorporation into muscle sarcomeres in cultured skeletal muscle myotubes formed by C2C12 cells**. **A**. Images of individual myofibrils within cultured skeletal muscle myotubes expressing eGFP–MHC constructs as shown. **B.** Example line profile for MYH7 WT from the image shown in panel A. Positions of the peak intensity values and the minimum intensity value (at the M-line) and ends of the sarcomere (at the Z-line) are indicated. **C**. Mean fluorescence intensity profiles for eGFP–MHC organization across a single sarcomere, for WT and each mutant. The mean plot profile was calculated from measurements of at least 30 sarcomeres from different myotubes from a minimum of three separate biological replicates. (n = 50, MYH7 WT; n = 35, A1440del; n = 53, L1492P; n=56, E1508del; n=81, E1610K; n=41, E1669del; n=30, K1729del; n=69, MYH2 WT). The MYH2 L1871P mutant did not express well enough to quantify sarcomere incorporation.

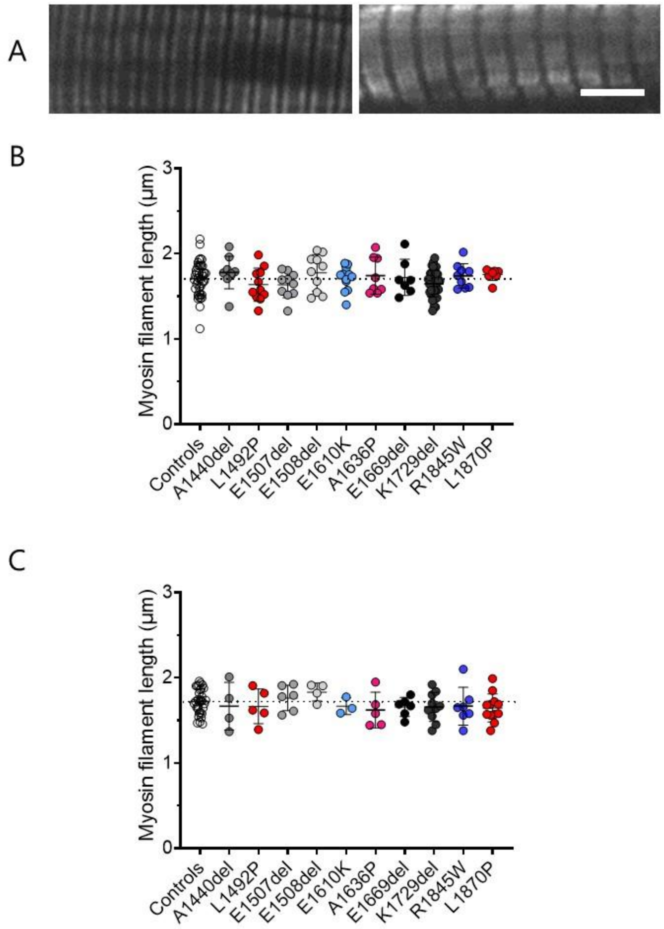


**Figure S3: Myosin filament length. A.** displays two typical images obtained with confocal microscopy using the A4.951 antibody and allowing the measurement of myosin filament lengths (scale bar: 5 µm). **B.** shows measurements for individual myofibres expressing the β/slow myosin heavy chain isoform in every single subject whilst **C** has data relating to muscle fibres expressing the type IIA myosin heavy chain isoform. Means and standard deviations also appear on these graphs.

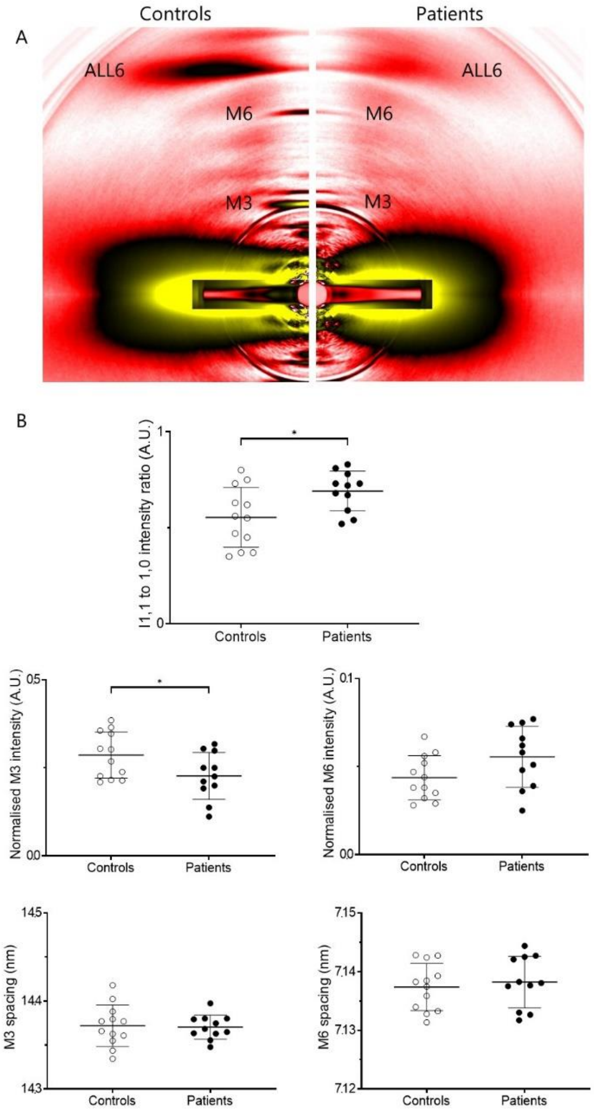


**Figure S4 Myosin head order. A.** depicts typical X-ray diffraction patterns. **B.** shows equatorial intensity ratio (IR) and the main myosin meridional reflections, namely M3 and M6 (M3 and M6 intensities were normalised to the $6^{th}$ actin-layer line, ALL6). To ensure reliable results and avoid misinterpretation, we pooled all the patients' data together and compared these to images acquired for controls. Means and standard deviations also appear on these graphs. $*p < 0.05$.

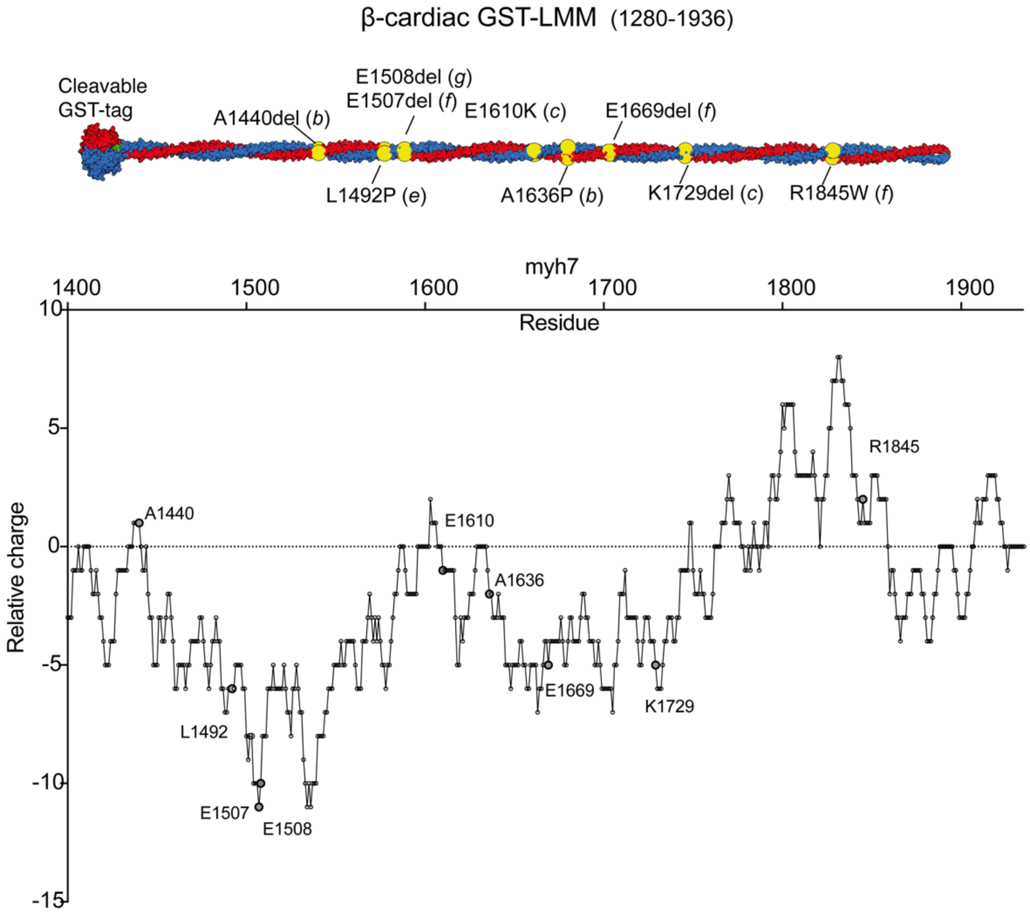


**Figure S5. Charge plot of MYH7 LMM region.** The diagram shows the positions of mutations in GST-LMM, and the plot shows the alternating regions of positive and negative charges important for filament formation, together with the positions of the mutations.
